# Turning the Fluorescent Protein Barrel into a Programmable Electrostatic Device Reveals Differential Bleaching States of the Chromophore

**DOI:** 10.64898/2026.02.12.705679

**Authors:** Nazarii Frankiv, Lee Min Leong, Bradley J. Baker

## Abstract

Genetically encoded voltage indicators (GEVIs) based on fluorescent proteins (FPs) report membrane potential changes through voltage-driven conformational rearrangements of a voltage-sensing domain that perturb the electrostatic and hydrogen-bonding environment of a fused FP chromophore. While the chromophore is largely protected from the external environment by the FP β-can, previous results suggest that this structure can also act as a programmable electrostatic grid that influences chromophore flexibility and fluorescence transition states. Here, we show that varying the external polar offset of β-sheet residues flanking a chromophore-proximal position systematically reshapes fluorescence transitions, altering response polarity and kinetics. Importantly, the influence of these external residues depends strongly on the internal sidechain chemistry, revealing reciprocal coupling between β-sheet electrostatics and chromophore-proximal residues. We further show that the functional effects of glutamine and asparagine substitutions at this position are strongly context-dependent and can be tuned by appropriate external polar offsets, consistent with electrostatic control of internal sidechain orientation and expanding the molecular switch repertoire. Disrupting steady-state fluorescence further revealed that the secondary component of the voltage-dependent fluorescence transition is preferentially light-sensitive and can be selectively diminished by repeated stimulation, indicating that it arises from a photophysically distinct process separable from the primary response. In addition, altering the chromophore protonation equilibrium (e.g., T65S) substantially changes response kinetics, indicating that chromophore state influences how β-can electrostatics couple to fluorescence transitions. Together, these results define general programming rules by which the electrostatic grid of the FP β-barrel can be tuned to control chromophore optical properties both at rest and during perturbation of the steady state.

## INTRODUCTION

Genetically encoded voltage indicators (GEVIs) enable optical readout of membrane potential by coupling voltage-driven conformational changes in a voltage-sensing domain (VSD) to fluorescence changes in a fused fluorescent protein (FP) domain[1, 2] (Figure 1A). Movement of the VSD in response to changes in membrane potential perturbs the electrostatic and hydrogen-bonding environment surrounding the FP chromophore, altering its photophysical properties and producing voltage-dependent fluorescence transitions[3]. Because the FP β-can structure largely isolates the chromophore from the external environment, these optical responses must arise from internal rearrangements within the FP domain[4], including changes in local electrostatics and sidechain interactions, that transmit voltage-induced conformational changes to the chromophore[5].

**Figure 1.**
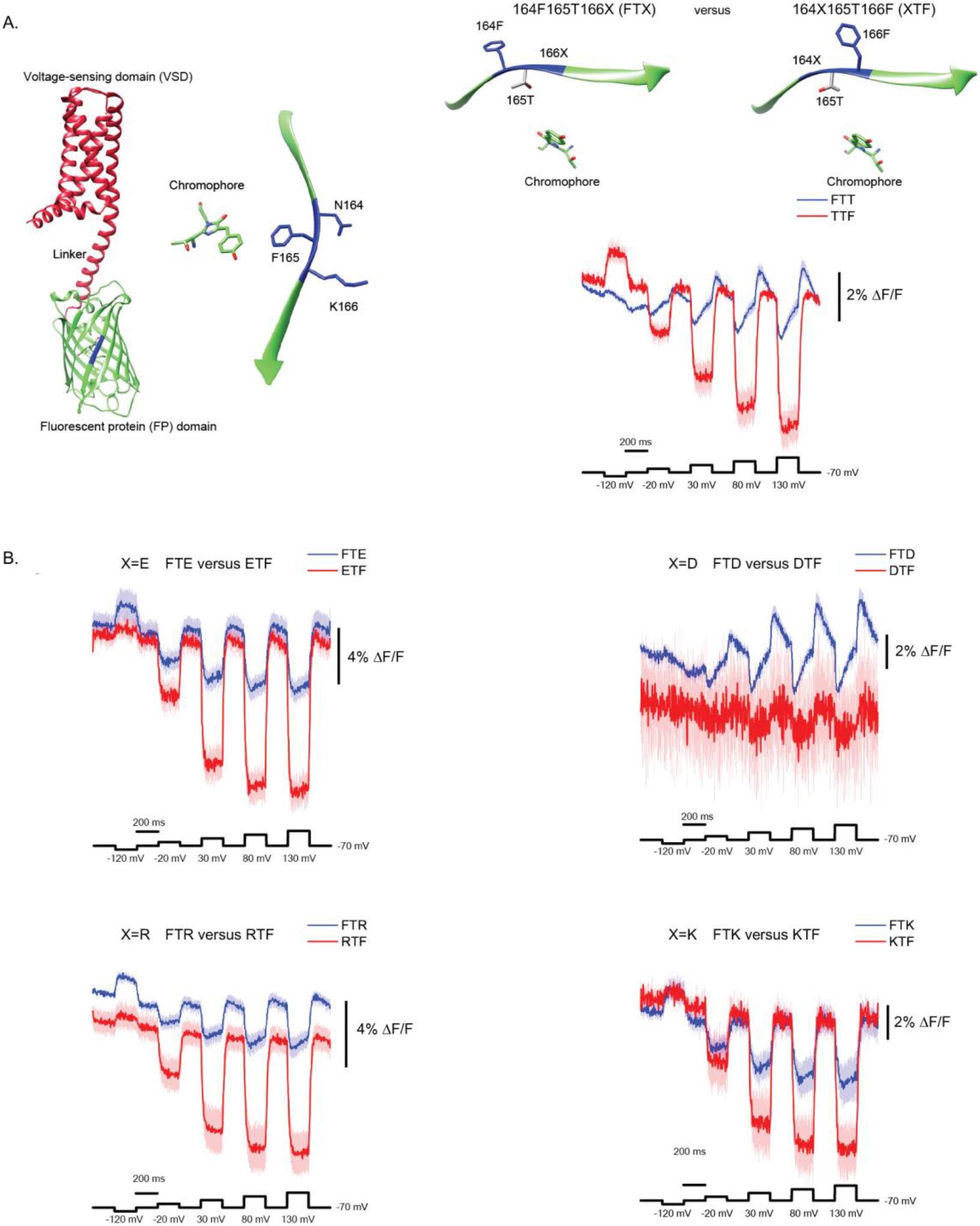
External β-strand electrostatics program voltage-dependent fluorescence transitions. A. Experimental design schematic. Left: AlphaFold3 [8] prediction of the TM GEVI, with the transmembrane voltage-sensing domain and cytoplasmic linker shown in red and the fluorescent protein (FP) domain in green. Middle: Zoomed view of the FP domain highlighting residues N164, F165, and K166 (blue). Right: Experimental design for altering the direction of the polar offsets across position 165, illustrating how external chemistry at positions 164 and 166 can bias the local electrostatic environment near the chromophore. Trace shows the directional effect of the external polar offset of 164F165T166T (blue trace) versus 164T165T166F (red trace). Structural figures made using Chimera[9]. B. Varying the external polar amino acid. Optical traces in response to voltage steps (black traces at bottom). Blue traces represent 164F165T166X. Red traces represent the opposite polar direction, 164X165T166F. Fluorescence signals were recorded from HEK293 cells under whole-cell voltage clamp at 1 kHz and low-pass filtered offline at 100 Hz. The dark center line is the mean of n number of cells. Shaded region represents the standard error of the mean. Directional changes in electrostatic context alter response polarity, kinetics, and the relative contribution of primary and secondary components.

Although the β-can provides a rigid protective scaffold for the chromophore, it is not an inert structure[6]. Local electrostatics, hydrogen-bonding networks, and sidechain interactions within the β-sheet framework can influence chromophore flexibility and fluorescence transition states. Structural and functional studies, including our analysis of the Ulla GEVI[3], have suggested that conformational changes can occur within internal regions of the FP domain during voltage sensing. These observations support the idea that the β-barrel can act as an electrostatically programmable environment rather than a purely insulating shell.

One chromophore-proximal element that strongly influences FP photophysics is the sidechain at position 165 (numbering relative to the FP domain), which lies near the chromophore and contributes to the local steric environment. In the wild-type FP, this position is occupied by phenylalanine, whose bulky aromatic sidechain primarily imposes steric constraints on the chromophore environment rather than participating in electrostatic interactions. Substitution of this residue with smaller or polar sidechains, such as alanine or threonine, reduces steric hindrance and enhances sensitivity to the electrostatic influence of neighboring β-strand residues, effectively coupling external β-sheet chemistry to the chromophore environment[3]. The orientation of an internal threonine sidechain at this position can be influenced by the chemistry of flanking β-strand residues, and such changes can strongly affect voltage-dependent fluorescence transitions. These findings suggest that external β-sheet residues can modulate internal sidechain behavior through electrostatic offsets, providing pathways by which VSD motion may be coupled to chromophore photophysics.

Importantly, deliberately inducing fluorescence transitions provides access to photophysical behaviors that are not observable under steady-state conditions. By driving the FP chromophore through repeated or prolonged voltage-dependent transitions, it becomes possible to reveal distinct light-sensitive states and differential bleaching behaviors of individual response components that report on underlying conformational and electrostatic rearrangements within the FP domain. Accordingly, voltage-driven fluorescence transitions provide a unique experimental handle for uncovering photophysical and conformational states of fluorescent proteins that are otherwise inaccessible.

These insights motivated us to systematically explore how electrostatic modulation extends across different sidechain chemistries at position 165 and how the internal steric context shapes sensitivity to external polar offsets. Our results demonstrate that steric constraint at this internal position strongly limits the influence of neighboring β-strand external residues, whereas permissive sidechains enable robust electrostatic tuning of fluorescence transitions. We further show that internal polar residues such as glutamine and asparagine exhibit context-dependent behavior that can be modulated by external β-strand chemistry, consistent with electrostatic control of internal sidechain orientation, representing an expansion of the molecular switch repertoire. Finally, we find that chromophore protonation state significantly alters the coupling between the β-can electrostatic grid and voltage-dependent fluorescence dynamics, revealing an additional layer of control over the biosensor’s optical responses.

Beyond guiding the design of FP-based biosensors, this type of targeted electrostatic interrogation provides a powerful experimental window into how protein β-strand chemistry and sidechain interactions shape internal conformational behavior. By linking external residue chemistry to internal sidechain orientation and photophysical outcomes, our approach reveals structural and electrostatic features of the FP β-barrel that are difficult to access through conventional structural methods alone.

## METHODS

### Plasmid Design and Construction

The triple mutant genetically encoded voltage indicator (GEVI) construct used in this study was based on the design previously described by Piao et al. (2015)[7], while the ArcLight configuration followed the description in Jin et al. (2012)[2]. Site-directed mutagenesis was performed to introduce specific point mutations within the fluorescent protein (FP) domain, using primers designed for each targeted residue. A two-step polymerase chain reaction (PCR) protocol was employed to generate the modified inserts. The resulting PCR products were digested with EcoRV and XhoI restriction enzymes (NEB) and subsequently ligated into the pcDNA3.1 expression vector (Invitrogen, USA) containing a cytomegalovirus (CMV) promoter to drive expression.

### Cell Culture and Transfection

HEK 293T cells were maintained in Dulbecco’s Modified Eagle Medium (DMEM; Gibco, USA) supplemented with 10% fetal bovine serum (FBS; Gibco, USA) at 37 °C in a humidified incubator with 5% CO_2_. For transfection, cells were detached using 0.25% trypsin-EDTA (Gibco, USA) and seeded onto poly-L-lysine–coated #0 glass coverslips (0.08–0.13 mm thickness, 10 mm diameter; Ted Pella, USA) at approximately 40–50% confluency.

Transfection was performed immediately after subculture to promote single-cell dispersion, enabling optimal conditions for patch-clamp recordings on the following day.

Transient transfection was conducted using Lipofectamine 2000 (Invitrogen, USA) according to a modified GenScript DNA transfection protocol.

### Electrophysiology

Coverslips containing transiently transfected HEK 293T cells were mounted in a patch chamber (Warner Instruments, USA) sealed with a #0 thickness glass coverslip to enable simultaneous voltage-clamp and fluorescence imaging. The chamber temperature was maintained at 34 °C and continuously perfused with extracellular solution consisting of 150 mM NaCl, 4 mM KCl, 1 mM MgCl_2_, 2 mM CaCl_2_, 5 mM D-glucose, and 5 mM HEPES (pH 7.4).

Patch pipettes were fabricated from filamented borosilicate glass capillaries (1.5 mm outer / 0.84 mm inner diameter; World Precision Instruments, USA) using a PC-100 micropipette puller (Narishige, Japan) to obtain tip resistances of 5–10 MΩ for HEK 293T cells. The pipettes were filled with an intracellular solution containing 120 mM K-gluconate, 4 mM NaCl, 4 mM MgCl_2_, 1 mM CaCl_2_, 10 mM EGTA, 3 mM Na_2_ATP, 5 mM HEPES, and 15 mM sucrose (pH 7.2).

Pipettes were secured in a pipette holder (HEKA, Germany) mounted on a micromanipulator (Scientifica, UK). Whole-cell voltage-clamp and current-clamp recordings were performed using a patch-clamp amplifier (HEKA, Germany).

### Fluorescence Microscopy of Cultured Cells

Epifluorescence imaging was conducted using an inverted microscope (IX71; Olympus, Japan) equipped with a 60× oil-immersion objective lens (numerical aperture, 1.35). Illumination was provided by a 470 nm LED light source driven by a DC4100 four-channel LED controller (ThorLabs, USA). GFP fluorescence was detected using a Semrock filter cube (USA) consisting of an excitation filter (FF02-472/30-25), a dichroic mirror (FF495-Di03), and an emission filter (FF01-496/LP-25).

Two cameras were mounted on the microscope via a dual-port camera adapter (Olympus, Japan). A low-speed color CCD camera (Hitachi, Japan) was used for cell visualization and patch positioning, while fluorescence recordings from voltage indicators were acquired at a 1 kHz frame rate using a high-speed CCD camera (RedShirtImaging, USA).

All optical components were placed on a vibration isolation platform (Kinetic Systems, USA) to minimize mechanical noise during patch-clamp fluorometry experiments.

### Optical Data Acquisition and Analysis

Images obtained from patch-clamp fluorometry were processed and analyzed to quantify fluorescence dynamics. Initial parameters were calculated as the fluorescence change (ΔF = F_x_− F_0_), where F_x_ represents the fluorescence intensity at frame x, and F_0_ is the average fluorescence of the first five frames of each recording. Fractional fluorescence changes were expressed as ΔF/F = ((F_x_ − F_0_) / F_0_) × 100. Data acquisition and initial analyses were performed using NeuroPlex (RedShirtImaging, USA) and Microsoft Excel (Microsoft, USA). Data from whole-cell voltage-clamp recordings of HEK 293T cells were analyzed as the average of four trials, each trial consisting of four repetitions (for a total of 16 technical replicates), unless otherwise noted. Only recordings from cells that maintained stable seals throughout the voltage-clamp protocol were included in the analysis. ΔF/F values corresponding to the applied voltage pulses were plotted using OriginPro 2016 (OriginLab, USA). For each recorded cell, ΔF/F traces over time were fitted with single- or double-exponential decay functions in OriginPro 2016.

## Results

### Directional electrostatic context at external positions 164–166 programs multi-component voltage responses

Previous work demonstrated that replacing the native phenylalanine at position 165 with threonine enables a secondary component in the voltage-dependent fluorescence response due to the loss of steric hindrance. However, sufficient steric hindrance could be functionally restored by biasing the orientation of the threonine methyl group via the introduction of a polar offset at positions 164 and 166. Indeed, T165 acted as a molecular switch, with the 164T/166F external offset suppressing the secondary signal while the inverse offset of 164F/166T maintained the secondary component (Figure 1A).

Here, we tested whether this electrostatic control of the threonine switch is retained when the external polar offset is altered. To do so, we systematically modified residues at position 164 in the presence of a 165T/166F background, and inverted the offset by maintaining 164F/165T while introducing polar residues at position 166 (Figure 1B).These new variants expand the range of external electrostatic environments acting on the threonine sidechain without altering the voltage-sensing domain or linker context.

Systematic scanning and inversion of residues 164 and 166 revealed that the direction of the polar offset continues to strongly influence voltage-dependent fluorescence (Figure 1B). Polar substitutions at position 166 promoted a pronounced secondary component, whereas polar substitutions at 164 suppressed it (an exception was 164D discussed below), indicating that the threonine switch remains electrostatically tunable across multiple chemical contexts. Inverting the 164–166 polarity arrangement altered response polarity, kinetics, and the relative contribution of the secondary component, confirming that directional electrostatic programming of the threonine switch is preserved across diverse external chemistries. The FP β-barrel environment can be reprogrammed to control multi-component fluorescence transitions.

### Steric context at position 165 gates electrostatic programming by β-strand residues

During the 164 scan in a 165T/166F background, we observed a striking discontinuity: 164D/165T/166F (DTF) exhibited severely reduced resting fluorescence and negligible voltage responses, whereas 164E/165T/166F (ETF) produced robust signals. Importantly, 164D was not generally deleterious to FP fluorescence, as 164D/165F/166F (DFF) was bright and exhibited robust voltage-dependent fluorescence transitions (Figure 2A).These comparisons indicate that the effect of residue 164 on both brightness and voltage-dependent signaling is strongly context-dependent, emerging prominently when residue 165 is sterically permissive (e.g., threonine) and becoming muted in a bulky 165 background (phenylalanine). Together, these results suggest that residues 164 and 166 can more effectively influence the chromophore environment when steric constraints at position 165 are reduced.

**Figure 2.**
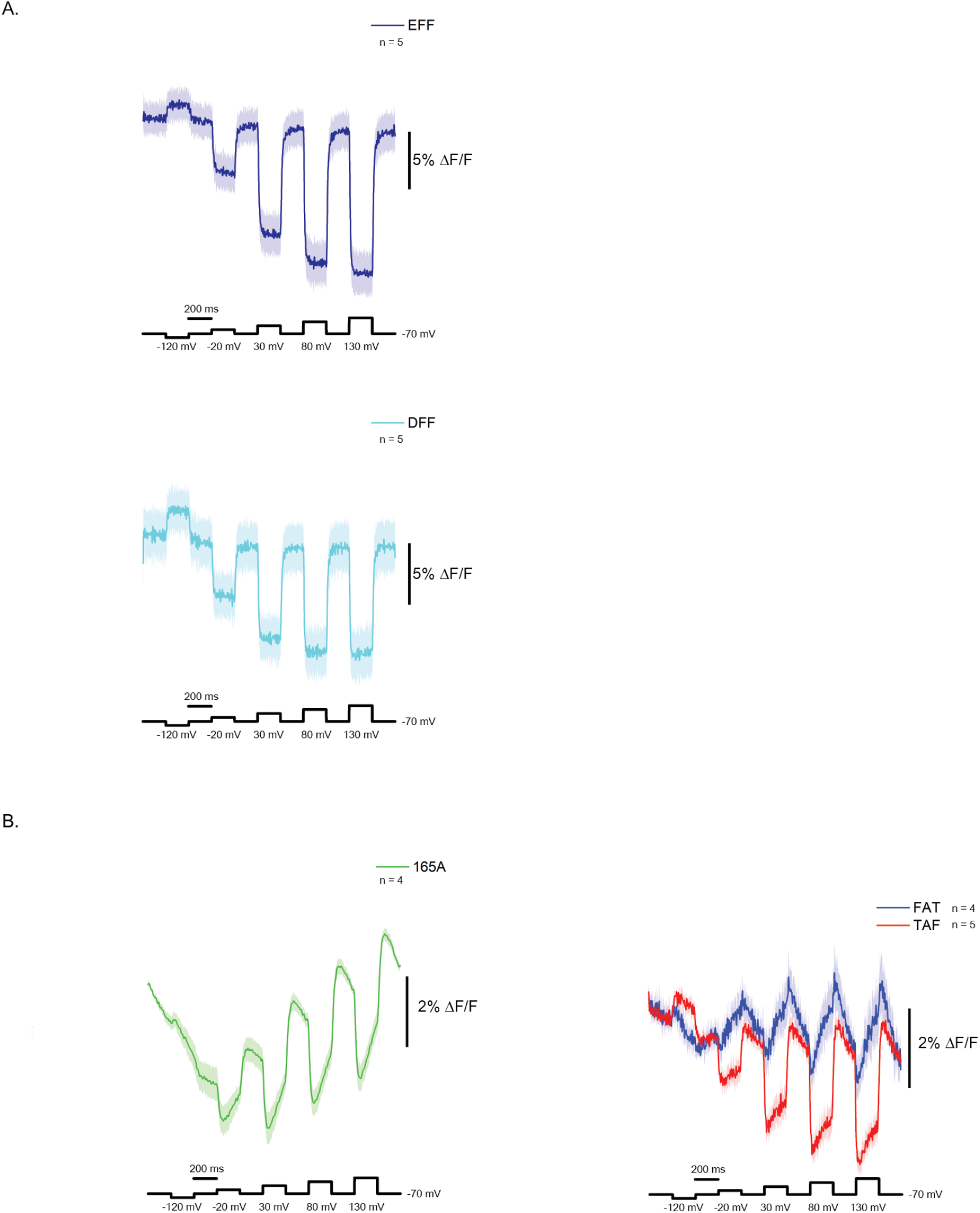
Steric context at position 165 gates electrostatic tunability by residue 164. A. In a bulky 165F/166F background, both acidic amino acids at position 164 produce a robust, voltage-dependent signal. B. In the sterically permissive background of 165A (green trace), external β-strand context produces larger changes in response shape and early kinetics depending on the orientation of the external polar offset (164F/165A/166T – blue trace, 164T/165A/166F – red trace) demonstrating that relief of steric constraint at position 165 unmasks strong electrostatic control of the fluorescence transition.

The 165A background provides a critical control because alanine lacks both steric bulk and a polar sidechain, and therefore should not exhibit orientation-dependent behavior. In this context, inversion of the 164–166 polar offset would ideally have little effect, serving as a measure of the intrinsic influence of external β-strand electrostatics in the absence of an orientable internal sidechain. Surprisingly, however, the 164F/165A/166T (FAT) and 164T/165A/166F (TAF) constructs exhibited distinct voltage-dependent responses, with marked differences in the relative weighting of the initial and secondary components (Figure 2B). This indicates that residues at positions 164 and 166 can directly reshape fluorescence transitions even when internal sidechain orientation is not a contributing variable. Importantly, these results demonstrate that the functional consequences of external β-strand electrostatics depend on the chemical context at position 165, suggesting that internal sidechain identity determines how external polar offsets are transduced into optical signals.

### External electrostatic modulation also influences the orientation of internal glutamine and asparagine sidechains: expansion of the molecular switch repertoire

Given that external β-strand electrostatics reshape voltage-dependent fluorescence in a manner conditioned by steric context at position 165, we next examined whether polar sidechains such as glutamine and asparagine could support distinct, context-dependent optical responses. Substitution of residue 165 with glutamine or asparagine alone (F165Q and F165N) yielded constructs with no detectable voltage-dependent fluorescence (Figure 3). Likewise, constructs containing these residues within a 164T/166F context (TQF and TNF) remained nonfunctional. In contrast, inversion of the external polar offset (FQT and FNT) restored resting fluorescence and produced measurable voltage-dependent signals. Similar to the 165A constructs (Figure 2), these rescued variants exhibited differences in the relative weighting of the initial and secondary components.However, whereas this weighting was determined by the orientation of the flanking polar offset in the 165A background, the relative contributions of the initial and secondary signals in the 165Q and 165N variants were dictated by the chemical identity of the residue at position 165.

**Figure 3.**
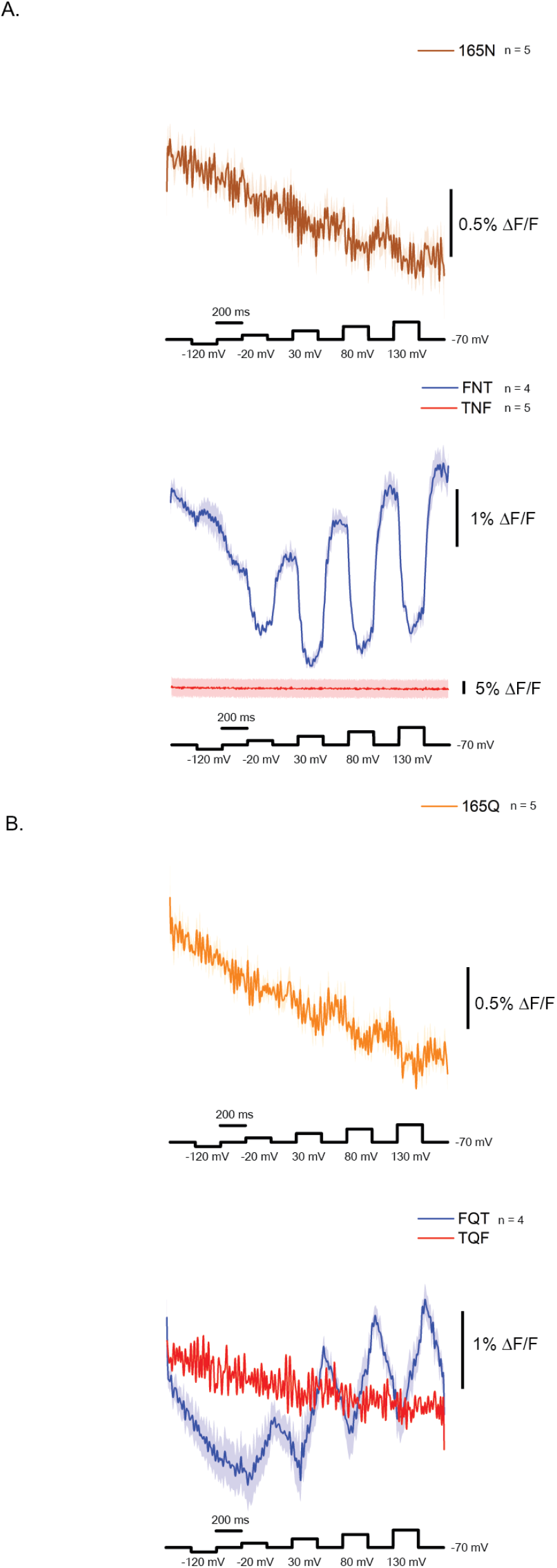
External β-strand chemistry enables context-dependent function of glutamine and asparagine at position 165. A. Substitution of residue 165 with glutamine (F165Q) yields dim constructs with negligible voltage-dependent fluorescence under baseline external contexts. Polar offset 164F/165Q/166T (blue trace) rescues voltage-dependent optical signal. Opposite polar offset (164T/165Q/166F – red trace) does not. B Substitution of residue 165 with asparagine (F165N) does not yield a voltage-dependent response. Polar offset 164F/165N/166T (blue trace) rescues voltage-dependent optical signal. Opposite polar offset (164T/165N/166F – red trace) does not.

### Disruption of steady state fluorescence reveals differential bleaching of the secondary component

One potential mechanism underlying the secondary component is that during the transition there is a reorientation of the chromophore within the FP barrel. The results in Figures 2 and 3 indicate that this secondary response represents a distinct and regulatable photophysical process whose contribution depends on both external β-strand electrostatics and chromophore-proximal sidechain chemistry. Notably, bulky sidechains at position 165 suppress the secondary component regardless of external polar offsets, whereas specific electrostatic contexts at positions 164 and 166 modulate its relative contribution when steric constraints at 165 are relieved. If the secondary component reflects a structural rearrangement such as chromophore rotation, it should exhibit measurable kinetics that can be probed by prolonging the depolarization. To test this possibility, we extended a single 200 mV depolarization step to 500 ms in order to drive the response toward a plateau and directly examine the temporal evolution of the secondary component.

The 164F/165T/166E (FTE) construct was expressed in HEK cells and submitted to the prolonged voltage transient. A representative example is shown in Figure 4A. The initial trial is shown in solid blue and the last trial is shown in cyan. Trials 2-15 are depicted in gray. Notice that the secondary signal is significantly more light-sensitive than the original voltage-dependent optical response. Figure 4 B compares the first trial to the last trial which shows a substantial decrease in the amplitude of the secondary response. Subtracting the light-exposed trial 16 from the initial trial yields the bleached secondary component. These data indicate that the secondary optical component is photophysically separable from the primary response and is preferentially light-sensitive. One plausible explanation is that the secondary component reflects a light-sensitive structural rearrangement within the FP barrel—potentially including chromophore reorientation—whose occupancy or reversibility is reduced by repeated illumination.

**Figure 4.**
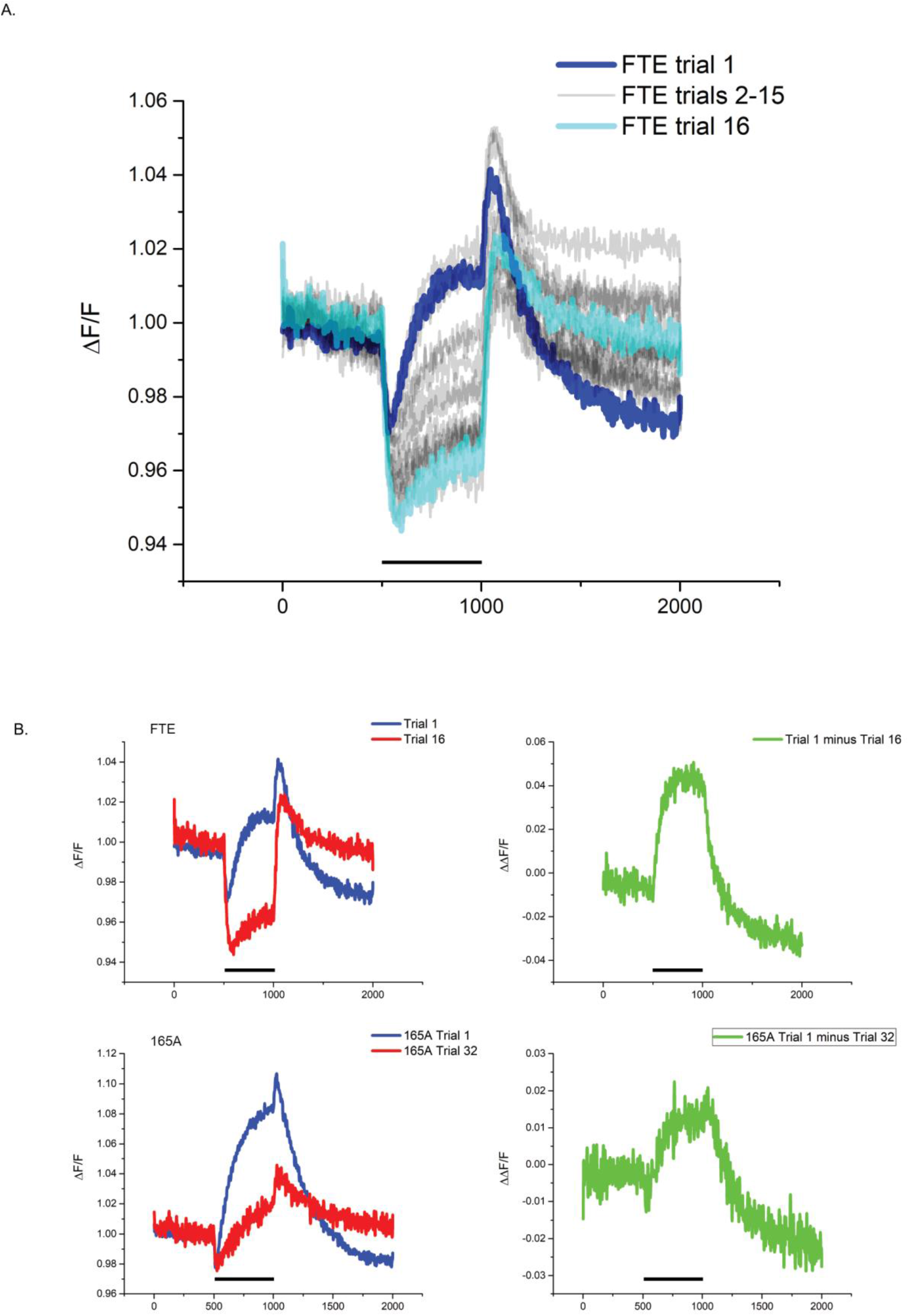
The secondary component is a separable, more light-sensitive component of the voltage-dependent response. A. An HEK cell expressing FTE was subjected to a 200 mV depolarization step for 500 ms and was repeated for a total of 16 trials. The holding potential was -70 mV. B. Left panel compares trial 1 (blue) to trial 16 (red) for both 164F/165T/166E (FTE - top traces) and 165A (lower traces). Right panel isolates the light-sensitive secondary component by subtracting trial 16 from trial 1. The black bar represents the time of the 200 mV step of the plasma membrane potential.

### Chromophore-state–dependent voltage coupling revealed by 390 vs 470 nm excitation

To further investigate the internal conditions resulting in the variable voltage-dependent optical response, we examined the potential role of protonated chromophore (excited at 390 nm) versus the role of the anionic chromophore (excited at 470 nm)[10, 11].Constructs such as FTE exhibited a prominent secondary component under 470 nm excitation, which preferentially reports on the anionic chromophore, whereas responses at 390 nm were smaller and often lacked a clear secondary phase (Figure 5). By comparison, ETF displayed a markedly reduced secondary component, highlighting the construct-specific nature of chromophore-state coupling.

**Figure 5.**
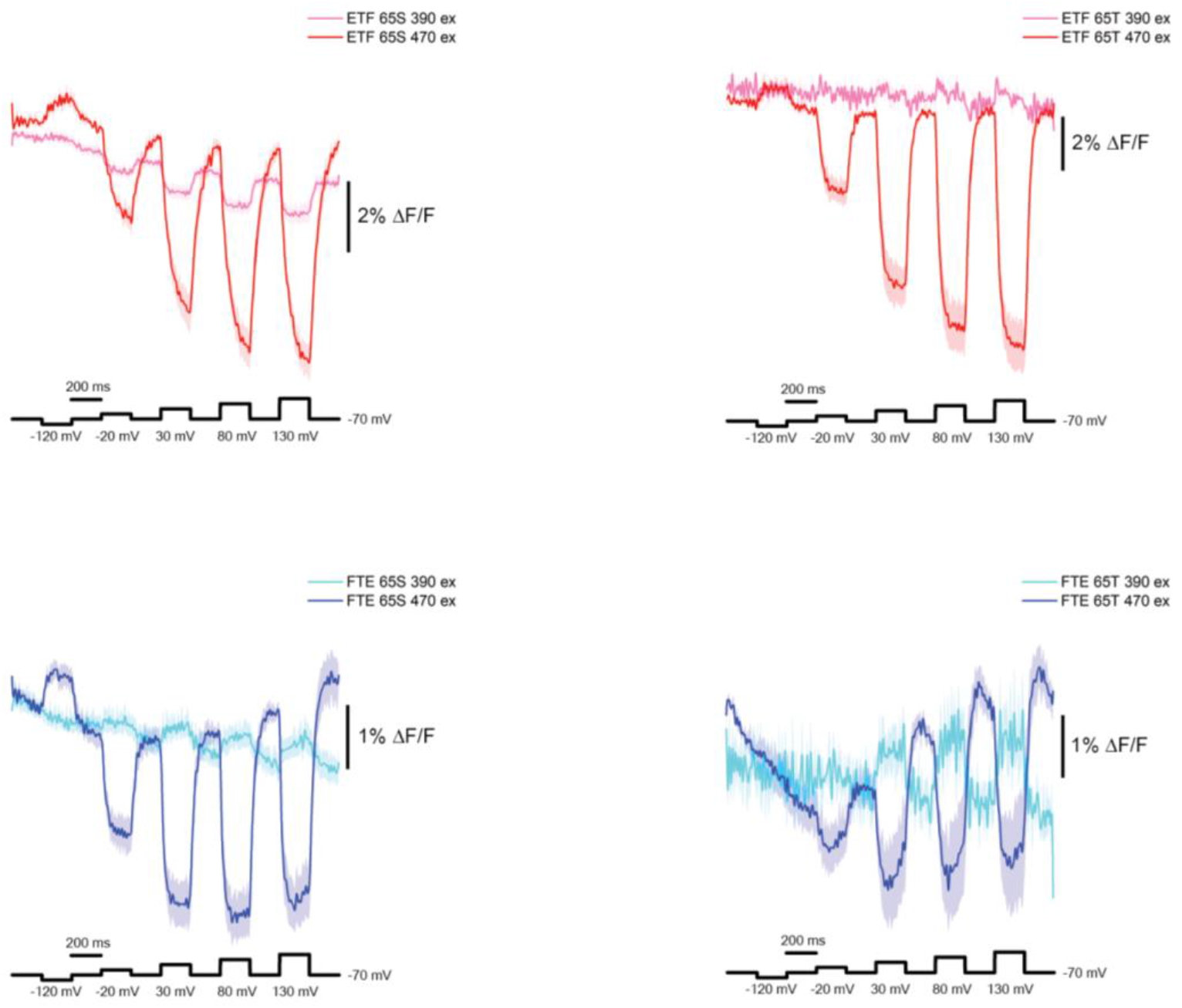
Chromophore protonation state modulates electrostatic coupling between the β-can and fluorescence dynamics. Excitation at 470 nm (anionic chromophore) reveals strong secondary components and faster kinetics, whereas 390 nm excitation (protonated chromophore) produces smaller and often slower responses. T65S variants exhibit altered kinetics even under 470 nm excitation, indicating that chromophore–barrel interactions shape voltage-dependent dynamics independently of chromophore charge state.

Notably, chromophore-state dependence extended beyond response amplitude and polarity to include response kinetics. Even when exciting the anionic chromophore at 470 nm, T65S variants displayed slower voltage-dependent kinetics than their T65 counterparts. This demonstrates that chromophore–barrel interactions influence voltage-dependent dynamics independently of chromophore charge state, further supporting the role of the FP-domain environment in shaping how voltage perturbations are transduced into fluorescence.

### Proton network and spatial manifestation

Mutation of Q149 to phenylalanine in a T65S background significantly enhanced voltage-dependent responses under 470 nm excitation while suppressing responses under 390 nm excitation, effectively restricting voltage sensitivity to the anionic excitation channel. This indicates that residues in the H148/Q149 region play a key role in determining which chromophore state participates in voltage reporting.

In addition to altering chromophore-state selectivity, proton-network mutations also reshaped the kinetics of the voltage-dependent response. In T65S/Q149F, responses recorded at 390 nm were faster than those recorded at 470 nm, despite the larger amplitude observed under 470 nm excitation.

The difference in kinetics had important consequences for spatial sensitivity. Curvature-dependent voltage responses were more readily observed under 390 nm excitation, where responses were fast, than under 470 nm excitation, where slower kinetics made curvature effects more difficult to resolve, even within the same cell.

Finally, mutation of H148 to threonine preserved the secondary voltage-dependent component and enhanced its spatial correlation with membrane curvature. Together, these results demonstrate that the chromophore proton-network environment not only determines chromophore-state selectivity and kinetics, but also shapes the spatial manifestation of voltage-dependent fluorescence signals.

The crystal structure of Super Ecliptic pHluorin, Ulla[3] and Lime[12] revealed the breaking point for the 7^th^ β-strand in the FP domain to occur at position 149. This unique structure for fluorescent proteins results in H148 being exposed to the external surface of the β-can structure. However, for Ulla at alkaline pH conditions, the β-strand is extended to the 148 position which orients H148 internally resulting in a hydrogen bond with the chromophore. To help determine the orientation of H148 in the GEVI expressed in the cell, we introduced the Q149F mutation placing a polar offset across H148 with D147 (147D/148H/149F).

Given that the protonation state of the chromophore has been shown to influence the kinetics and polarity of the voltage-dependent optical signal[5], we also introduced the T65S mutation to better excite the protonated chromophore with 390 nm illumination. Figure 6 reveals an interesting characteristic of this probe. First is that the kinetics of the anionic chromophore are much slower than the protonated chromophore. Second is that the protonated chromophore exhibits a noticeable secondary component during the fluorescence transition that is influenced by the curvature of the plasma membrane. Voltage transients during 470 nm illumination of the same cell also exhibits a slight curvature effect but is much diminished. These results suggest that the flexibility (secondary component) is higher in the protonated chromophore and that the anionic chromophore is much slower to respond (slower initial component).

**Figure 6.**
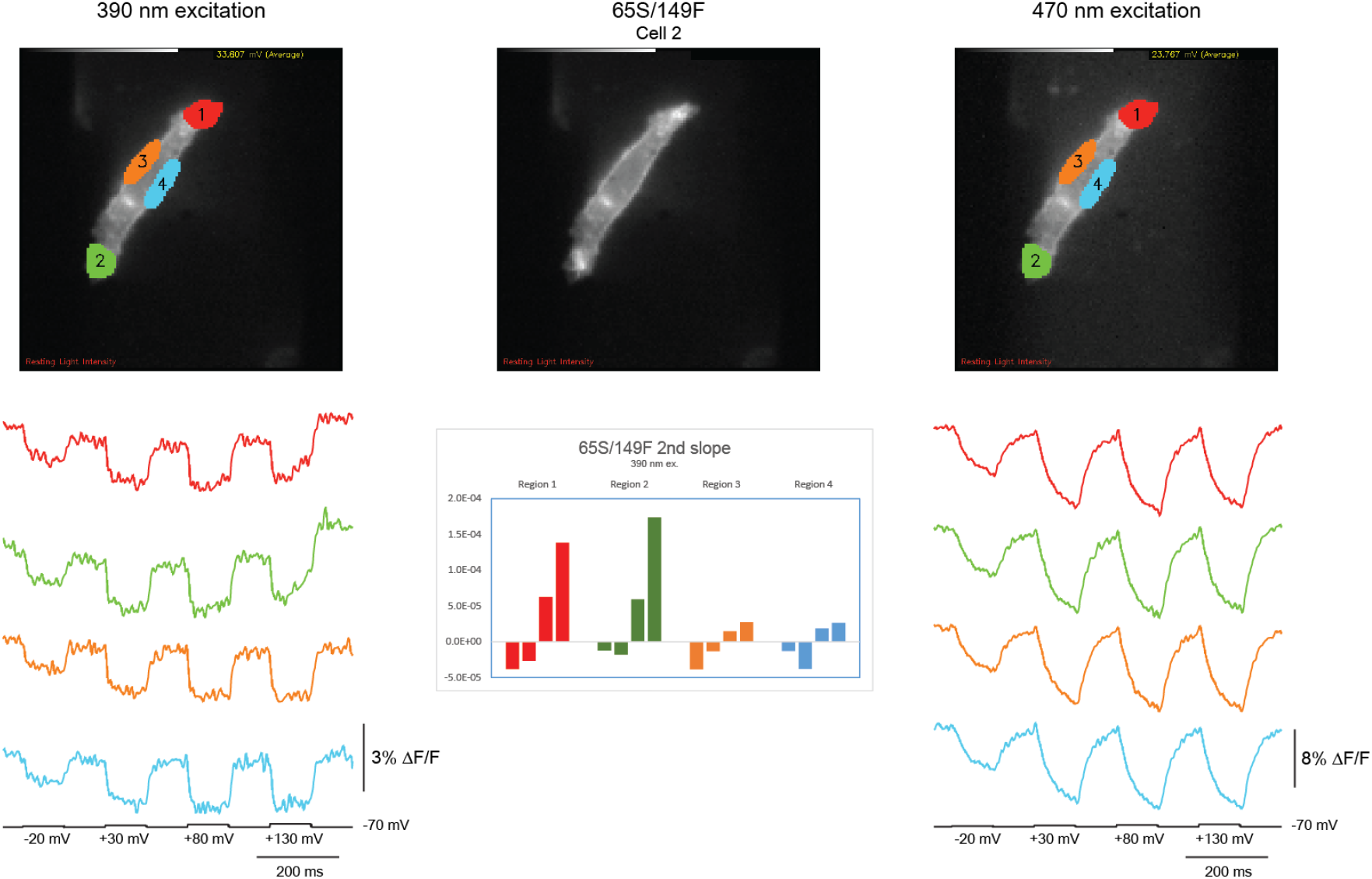
Chromophore protonation state reveals curvature effect in the 65S/149F context. Introducing the Q149F mutation expands the polar offset strategy (D147/H148/F149) to other regions of the FP domain. Representative cell with S65 mutation to improve 390 excitation and F149 to introduce a novel polar offset yields a GEVI that at 390 excitation exhibits an obvious curvature effect on voltage-dependent optical responses while excitation at 470 nm exhibits a much reduced curvature effect.

## DISCUSSION

In this study, we demonstrate that the fluorescent protein β-barrel is not merely a passive scaffold that insulates the chromophore, but instead functions as a programmable electrostatic environment capable of shaping voltage-dependent fluorescence transitions. By systematically manipulating the external polar context of β-strand residues flanking a chromophore-proximal position, we show that internal sidechain orientation, chromophore coupling, and fluorescence kinetics can be modulated (Figure 2).

A central finding is that electrostatic programming is strongly gated by steric context at position 165 (Figure 2). When steric constraints near the chromophore are high, external β-strand substitutions exert limited influence on the optical response. In contrast, sterically permissive sidechains allow residues at positions 164 and 166 to bias fluorescence transitions through directional electrostatic offsets. This epistatic relationship reveals how local sterics and electrostatics jointly determine the functional landscape of the FP domain.

Importantly, this electrostatic control is not restricted to threonine (Figure 3). We find that internal polar residues such as glutamine and asparagine, which are nonfunctional in baseline contexts, can be “licensed” to produce voltage-dependent signals through appropriate external β-strand chemistry. This expands the repertoire of chromophore-proximal residues whose behavior can be tuned, underscoring the generality of electrostatic programming within the FP β-barrel.

Disrupting the fluorescence steady state further revealed that voltage-dependent responses comprise multiple photophysical components with distinct sensitivities to illumination (Figure 4)[13, 14]. The preferential bleaching of the secondary component indicates that it arises from a light-sensitive process separable from the primary response, highlighting the dynamic nature of FP photophysics under voltage perturbation. These observations suggest that GEVI experiments provide access to chromophore behaviors that are difficult to resolve using static fluorescence assays alone.

A central implication of the differential bleaching behavior is that the voltage-dependent fluorescence transition comprises at least two photophysical processes with distinct photostability. In these GEVIs, voltage does not act directly on the chromophore; instead, voltage-driven motion of the VSD is transduced through changes in FP-domain interactions (e.g., altered FP–FP packing or interface geometry), which then reshapes the internal chromophore environment. One mechanistic possibility is that the secondary component reflects an internal FP rearrangement—potentially including chromophore reorientation or coupled β-strand remodeling—that is promoted by a particular FP–FP interaction state and is preferentially depleted or trapped by illumination. Alternative explanations include light-dependent remodeling of hydrogen-bond networks or local β-strand flexibility near the chromophore. Distinguishing among these models will require experiments that constrain interdomain geometry and/or directly report on barrel dynamics.

In addition to revealing separable photophysical processes (Figure 4), our results demonstrate that chromophore-state identity strongly influences how β-barrel electrostatics are transduced into optical responses (Figure 5). Excitation-selective probing of the anionic and protonated chromophore populations revealed distinct kinetic behaviors and differential engagement of the secondary component, indicating that electrostatic programming does not act uniformly across chromophore states. Rather, the β-barrel electrostatic grid modulates chromophore dynamics in a state-dependent manner, providing an additional layer of tunability in voltage-dependent fluorescence transitions. Together with the steric and chemical gating observed at position 165, these findings establish a hierarchical framework in which interdomain coupling, local electrostatics, and chromophore protonation state collectively define the optical output of the FP domain.

The effects of Q149F further suggest that electrostatic programming within the FP barrel may extend beyond residue 165 to structurally dynamic regions of the β-sheet framework. Because residue 149 lies at a known discontinuity in the seventh β-strand, mutations at this position have the potential to alter strand extension and local packing, thereby influencing the positioning of neighboring residues such as H148. The chromophore-state–specific effects observed in the 149F background are consistent with a model in which β-strand register or hydrogen-bonding interactions near the chromophore are dynamically modulated during voltage coupling. While the precise orientation of H148 in the cellular context remains to be determined, these findings underscore the structural sensitivity of the chromophore environment to subtle β-strand perturbations.

Together, these results define general programming rules by which the electrostatic grid of the FP β-barrel can be tuned to control chromophore behavior both at rest and during perturbation of the steady state. Beyond guiding the rational design of improved genetically encoded voltage indicators, this work illustrates how targeted electrostatic interrogation of β-strand chemistry can reveal internal conformational and photophysical properties of proteins that are otherwise challenging to access. In this way, voltage imaging platforms serve not only as sensors of cellular physiology, but also as experimental tools for probing fundamental principles of protein structure and dynamics.

